# Renal adaptation to high salt diet requires tubular Na^+^ secretion through type A intercalated cells

**DOI:** 10.64898/2026.02.02.702743

**Authors:** Chloé Rafael, Luciana Morla, Justine Billiet, Lydie Cheval, Samia Lasaad, Sandrine Placier, Christine Walter, Nicolas Picard, Gilles Crambert

## Abstract

**Background:** In the context of increased salt intake in the world population, the understanding of the mechanisms that contribute to its correct renal excretion and therefore, avoid variation of blood volume and blood pressure is of major importance.

**Methods:** Molecular, *ex vivo* microperfusion on isolated tubules, and integrative analysis, was used to identify, characterize and investigate a Na^+^ secretion pathway in the collecting duct.

**Results:** In collecting duct of mice, salt load induced an increase of the type A intercalated cells (AIC) number, an overexpression of the H(Na),K-ATPase type 2 (HKA2) catalytic subunit *Atp12a* and a stimulation of the bumetanide-sensitive Na^+^ secretion in isolated and microperfused tubules. Surprisingly, HKA2KO mice fed a high-salt diet exhibit a strong dysregulation of their Na^+^ and water balance with a pronounced loss of Na^+^ and fluid, alkalosis, hypokalemia and low blood pressure. This Bartter-like phenotype is due to an over-inhibition of the thick ascending limb (TAL) related to an elevated PGE2 production.

**Conclusion:** Our findings establish that activation of Na^+^ secretion in AIC act as the fine-tuning knob in the regulation of renal Na^+^ excretion in response to high salt intake. Its absence is overcompensated by an inhibition of the Na^+^ transport system of the TAL.

## Introduction

The modern Western diet affects electrolyte balance, producing approximatively 50 mmol of acid/day in contrast to our ancestors who were considered to be net base producers^1^. Additionally, the consumption of salt (NaCl) and potassium (K^+^) has shifted from a rich-K^+^/low NaCl diet in hunter-gatherer populations to the opposite in the modern, westernized population^2^. Our kidneys help us cope with these dietary changes, but long-term modifications can contribute to diseases like hypertension, especially with certain genetic polymorphisms that would have remained silent under more “natural” diet. All segments of the nephron contribute to salt balance, but the distal part (collecting duct, CD) is crucial for the fine regulation of Na^+^ and Cl^−^ excretion. The CD consists of principal cells (PC) and A– and B-intercalated cells (AIC and BIC). Until recently, these cell types were assigned to very specific functions i.e. the reabsorption of water and Na^+^ and the “normal” K^+^ secretion was attributed to PC whereas BIC were thought to be involved in base secretion in response to alkalosis and AIC in the H^+^ secretion in response to an acid load and both K^+^ reabsorption (for review see^3^) and K^+^ secretion^4^. This dogma is being revisited due to very recent observations demonstrating that BIC and AIC also contribute to salt balance by reabsorbing or secreting Na^+^ and Cl^−^, respectively^5^. The CD is divided into cortical and medullary parts; the latter lacks BICs, so only PC reabsorption and AIC secretion of Na^+^ influence the final urinary Na^+^ content. This study demonstrates that adaptation to a high-salt diet relies not only on inhibiting PC-dependent reabsorption but also on activating AIC-dependent Na^+^ secretion.

## Material and Methods

All material submitted conforms with good publishing practice in physiology^6^. We adhere to the NIH Guide for the Care and Use of Laboratory Animals or the equivalent. All animal experimentations were conducted in accordance with the institutional guidelines and the recommendations for the care and use of laboratory animals put forward by the Directive 2010/63/EU revising Directive 86/609/EEC on the protection of animals used for scientific purposes (project has been approved by a user establishment’s ethics committee and the Project Authorization: number 21927). Further details regarding the materials and procedures, are provided in the Supplemental Material and Methods.

## Animals

Experiments were performed on male C57BL/6JRj wild-type or knock-out mice for the *Atp12a* gene (HKA2KO)^7^. HKA2KO mice are regularly (every 5-10 generations) backcrossed with wild-type C57BL/6JRj male mice from the Janvier Labs to avoid genetic deviation. The animals were kept at CEF (Centre d’Explorations Fonctionnelles of the Cordeliers Research Center, Agreement no. A75-06-12).

## Metabolic analysis

To record physiological parameters, mice were placed in metabolic cages (Techniplast, France) for 2 days of adaptation and were fed a standard laboratory diet (NS, 0.8 % Na Cl; UPAE, INRA, Jouy-en-Josas, France) for at least two days and a high salt diet (HS, 8% NaCl; UPAE, INRA, Jouy-en-Jossas, France) for 3 to 6 days depending on the performed analysis. For GFR, urine and plasma parameters measurements see the Supplemental Material and Methods

Furosemide, thiazide and amiloride-sensitive natriuresis were measured as previously described^8,9^. To test for the vasopressin response, ddAVP (1.5µg/kg in 100µl of 0.9% NaCl) or its vehicle were injected intraperitoneally and a spot of urine was taken 5h post-injection for measurement of osmolality (Vogel 6300 Voebling osmometer, Bioblock Scientific) and compared to the osmolality of urine taken before injections.

Blood pressure was measured in conscious restrained mice by a tail-cuffed plethysmography method (BP2000, Visitech system, France) after a week of adaptation to the apparatus between 9:00 and 11:00 AM.

## RNA sequencing

RNAs were extracted from about 100-120 OMCD segments isolated from Liberase-treated kidney using RNeasy micro Kit (Qiagen). The quality of the extracted RNAs was checked using a bioanalyser 2100 (Agilent Technologies) and considered of good quality for a RIN > 7. The samples were then sent to the facility iGenSeq of the Institut du Cerveau et de la Moelle Epinière (Hôpital de la Pitié Salepêtrière, Paris, France) and processed for RNA sequencing using the Stranded mRNA kit (Illumina). The results have been deposited on the GEO website of the NCBI (GSE244598).

## AIC isolation

Kidneys from C57Bl6JRj mice were prepared as outlined in Picard et al.^10^ with some adaptations and AIC was sorted as single cells, as described in Chen et al.^11^ (see Supplemental Material and Methods for details).

## Sodium flux in isolated microperfused OMCD

Wild-type mice OMCDs were microdissected from the outer strip of the medulla and microperfused under symmetrical conditions. The bath and perfusate contained 118 mM NaCl, 23 mM NaHCO_3_, 1.2 mM MgSO_4_, 2 mM K_2_HPO_4_, 2 mM calcium lactate, 1 mM sodium citrate, 5.5 mM glucose, and 12 mM creatinine, at pH 7.4. The bath was continuously gassed with 95% O_2_/5% CO_2_ and heated at 37°C. Tubules were dissected for less than 60 min before mounting. After 30 min of equilibrium, 3 collections of 7 min were taken for the control period. Bumetanide 10^−4^M was added to the bath (basolateral side) and amiloride 10^−5^M or bafilomycine (4×10^−8^M) were added in the perfusion (apical side). After 30 minutes, 4 collections of 7 min were taken for the experimental period. The collection volumes were determined under water-saturated mineral oil with calibrated volumetric pipettes. Further details are provided in the Supplemental Material and Methods.

## Cell counting on isolated tubules

15-20 short OMCD segments were isolated from Liberase-treated kidney and transferred onto Superfrost Plus glass slides and treated as described recently in^12,13^. AIC were identified as AE1+ cells after labeling with an anti-AE1 antibody (1/500, gift from C.A. Wagner). For cell and doublet counting, a 3D reconstruction from all stacks (ImageJ) was performed from images of AE1-labelled OMCDs acquired by confocal microscopy^12,13^.

## Results

### High salt diet modifies gene expression of medullary CD

We conducted a RNA-seq analysis on isolated OMCD to identify genes implicated in the adaptation to a high-salt diet (3 days) in collecting duct of control mice (n=4). We quantified gene expression during a high-salt diet (HS, 8% NaCl) relative to a control diet (NS, 0.8% NaCl). As shown in Figure 1A and B, a total of 11688 genes were identified from which 293 are significantly differentially expressed (p<0.05). Among these genes, 117 genes were upregulated and 71 downregulated by HS (>50% compared to NS). Expression of genes involved in Na^+^ transport like *Scnn1a* (ENaC α subunit), *Scnn1b* (ENaC β subunit), *Sgk1* or in water transport like the water channels *Aqp2* and *Aqp3* are modified as expected by HS, which is confirmed by qPCR (Supplemental Figure 1). Gene expression analysis of GO gene sets enrichment highlighted the biological processes involved in the adaptative response to a high-salt diet. It turns out that the most affected processes are linked to cell proliferation/division, (Figure 1C and D).

**Figure 1:**
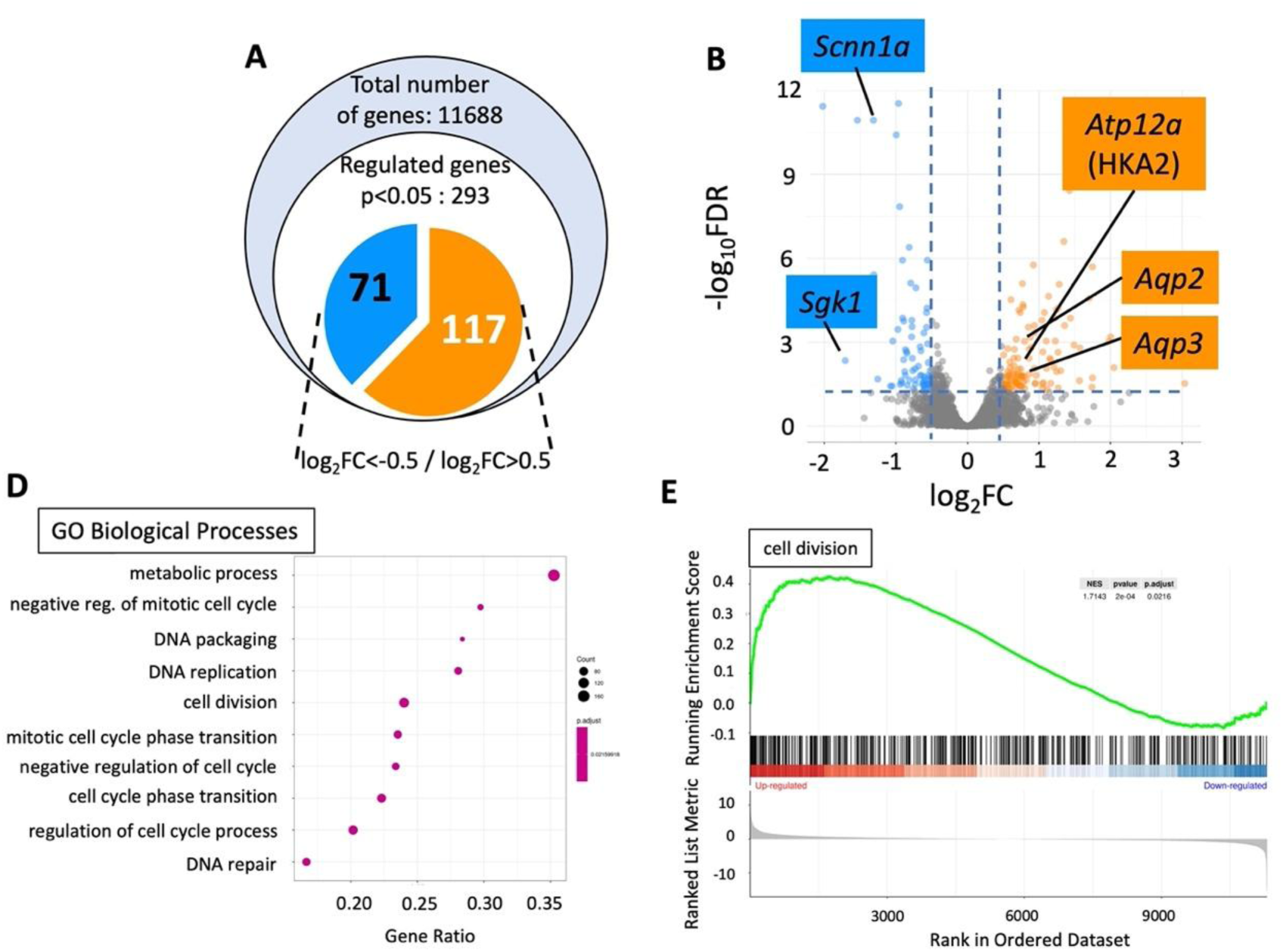
High salt diet modifies gene expression in collecting ducts of wild-type mice. **A)** Summary of the number of expressed genes that have been sequenced, found significantly regulated with a log2 fold changes (log2 FC) below –0.5 and above 0.5. **B**) Volcano plot highlighting Differentially Expressed Genes (DEG) at FDR<0.05 with log2 FC below –0.5 and above 0.5. **D-E**) Overexpression analysis of gene ontology dataset of biological processes and Gene set enrichment analysis (GSEA) of the GO Biological Process “cell division” showing significant positive enrichment among up-regulated genes (nominal p = 2 × 10⁻⁴, FDR-adjusted p = 0.0216), indicating activation of cell division–related processes in the analyzed condition.

### High salt diet increases number of AIC

Since the expression of genes involved in cell division were modified by HS, we quantified the cell composition of isolated OMCD by specifically labelling AIC with anti-anion exchanger 1 (AE1) antibody and nucleus with DAPI (Figure 2A). As shown in Figure 2B, the total number of cell/mm is increased by around 10% which may suggest an elongation of the medulla. More specifically, the proportion of AIC (AE1+ cells) is increased by 13% in OMCD from mice fed a HS (Figure 2C). The number of AE1-cells was not significantly increased by the HS diet (Figure 2D). The expression of a proliferative marker, *Pcna*, is significantly increased in isolated OMCD (Figure 2E) and *Pcna* expression exhibited a tendency to increase in AIC (p=0.09, Figure 2F).

**Figure 2:**
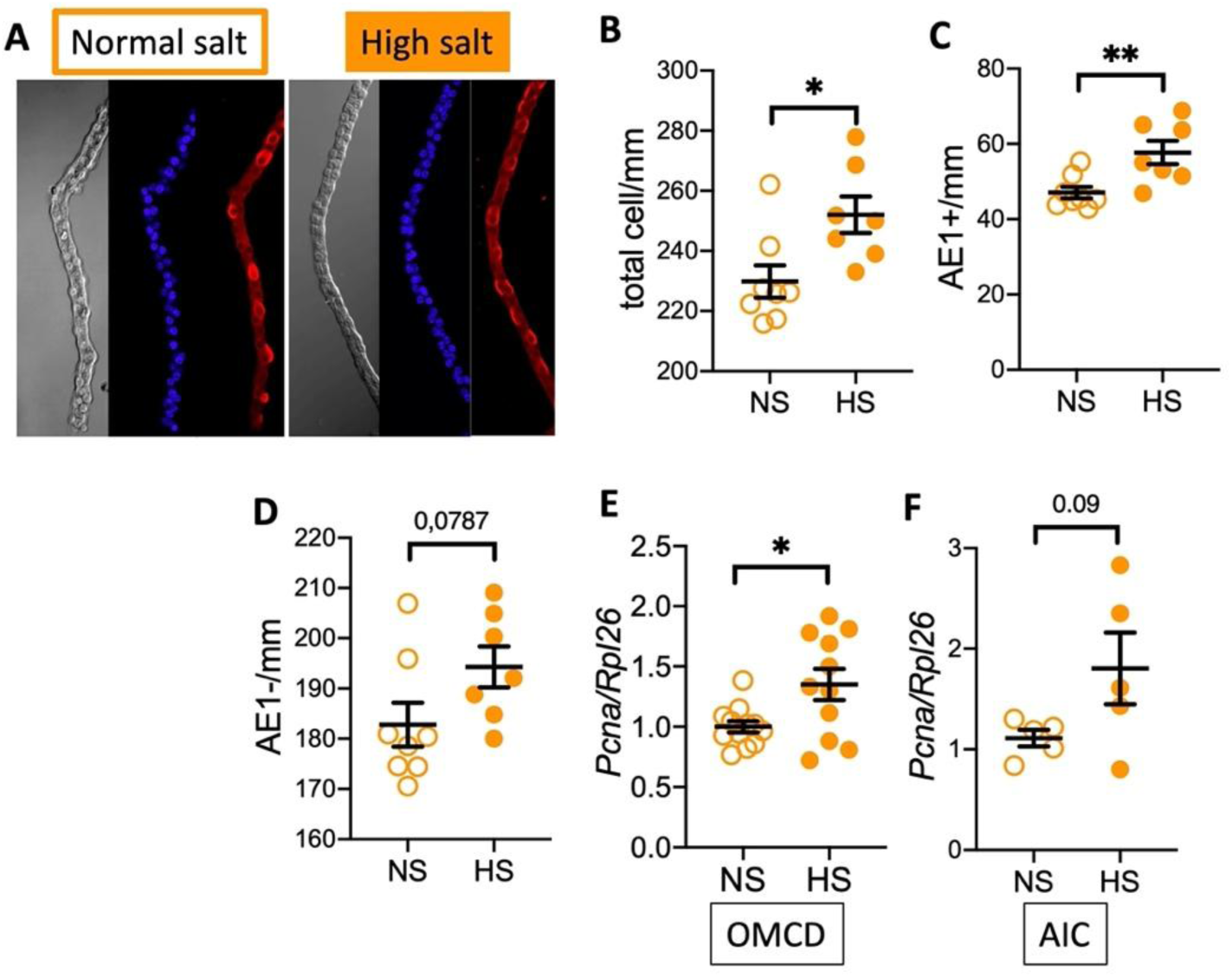
The number of ICA is increased by salt load. **A**) Examples of light and immunofluorescence confocal microscopy from isolated OMCD from WT mice under normal (NS) or high salt (HS) diets after labeling with DAPI (blue) and anti-AE1 antibody (red). The stacks of these images were then used to reconstruct tubules in 3D in order to accurately count the number of nucleus (**B**), of AE1+ cells (**C**) and AE1-cells (**D**) normalized by the length of the tubules. Each symbol represents the mean value of 7-11 reconstructed tubules of the same animal. Results are shown as mean±s.e.m (two series, n=8 under normal salt diet and n=7 under high salt diet) and analyzed by an unpaired Student t-test (*p<0.05, ** p<0.01). **E-F**) *Pcna* expression in isolated OMCD (n=11) and in isolated AIC (n=5) normalized by the housekeeping gene *Rpl26* in WT mice under NS or HS diets. Results are shown as mean±s.e.m and analyzed by an unpaired Student t-test (*p<0.05, ** p<0.01).

### High salt diet activates the secretion of Na^+^

In the RNA-seq analysis, we identified *Atp12a* encoding the H^+^-K^+^-ATPase type 2 (HKA2) catalytic subunit as one of the up-regulated genes by HS. We confirmed that *Atp12a* expression is 3-fold higher in microdissected medullary collecting ducts from mice under HS compared to NS (Figure 3A). We also showed that expression of *Atp12a* was strongly stimulated by HS in isolated A-type intercalated cells (AIC) (Figure 3B). We previously described the presence of a Na^+^ secretion pathway across AIC^14^ that is regulated by ANP and cGMP^15^ (Figure 3C). We observed that both plasma ANP and urine cGMP levels were increased in HS condition (Figure 3D-E).

**Figure 3:**
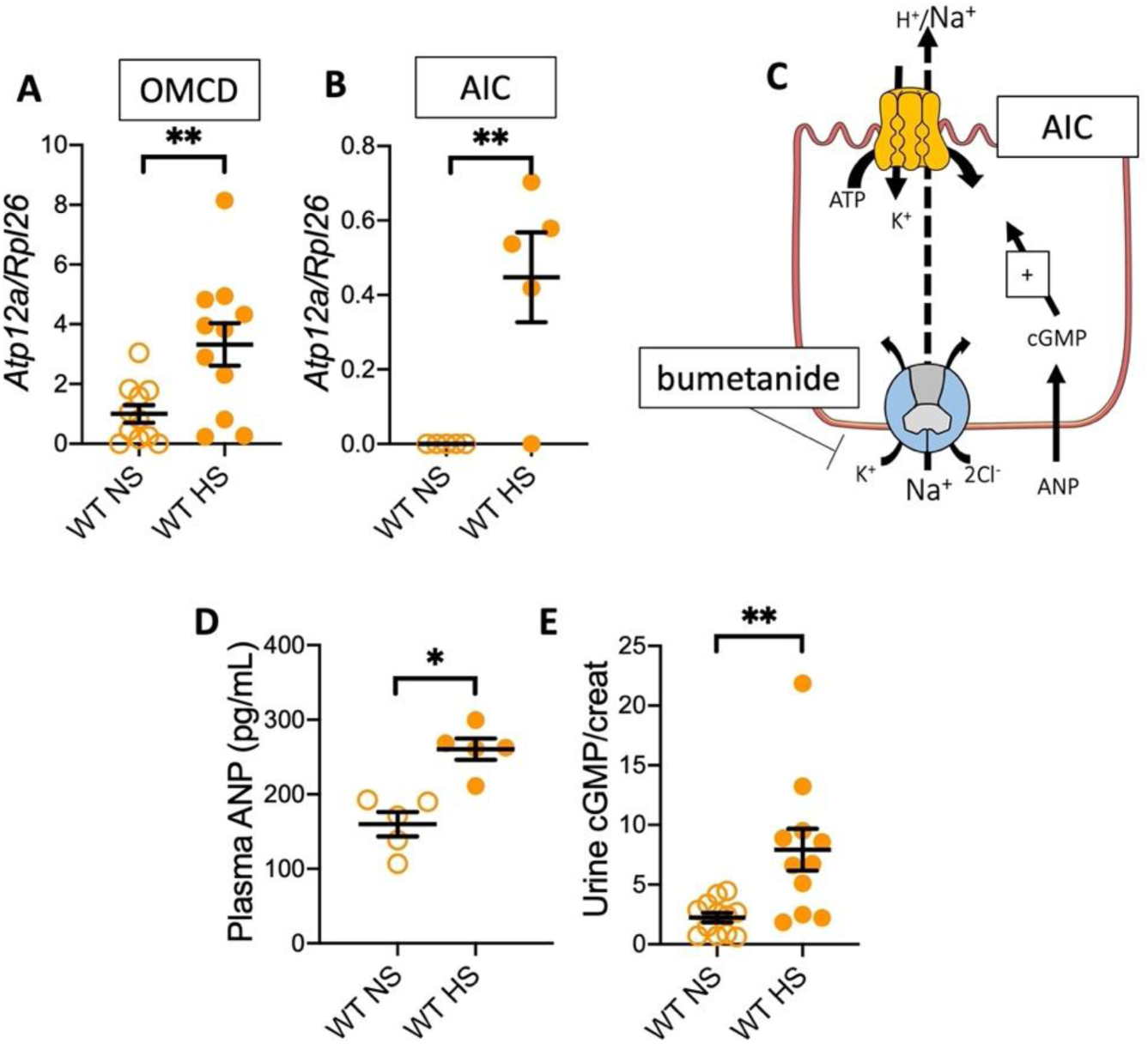
The secretion pathway of Na^+^ is upregulated by salt loading. **A-B)** *Atp12a* expression in isolated OMCD (n=11) and in isolated AIC (n=5) normalized by the housekeeping gene *Rpl26* in WT mice under NS or HS diets. Results are shown as mean±s.e.m and analyzed by an unpaired Student t-test (** p<0.01). **C**) Schematic representation of the secretion of Na^+^ in the AIC. **D**) Plasma concentration of the atrial natriuretic peptide (ANP) in WT mice under NS (n=5) or HS (n=4) diets. Results are shown as mean±s.e.m and analyzed by an unpaired Student t-test (* p<0.05). **E**) Urine excretion of cGMP normalized by the excretion of the creatinine on 24h collection in WT mice under NS or HS diets (n=11). Results are shown as mean±s.e.m and analyzed by an unpaired Student t-test (** p<0.01).

Microperfusion analysis allows to measure transepithelial ion fluxes that are the results of trans and paracellular reabsorption (positive values) and transcellular secretion (negative values) pathways (Figure 4A). We showed by microperfusion on isolated OMCD, that the transepithelial flux of Na^+^ (JNa^+^) was not affected by the presence of bumetanide (an inhibitor of NKCC1) at the basolateral side of the tubular cells in NS condition (Figure 4B-C). On the contrary, after HS period, the inhibition of NKCC1 led to an increase of the apparent reabsorption of Na^+^ (Figure 4B-C). The transepithelial Na^+^ flux results of the addition of cellular reabsorption (through PC) and secretion (through AIC) processes. A significant modification in JNa^+^ could therefore results either to modification of both processes. Here, the global increase of JNa^+^ is more likely due to an inhibition of a secretion process by bumetanide than an activation of reabsorption in the PC since we showed a reduction of ENaC expression (Figure 1 and Supplemental Fig 2) and that the context of HS diet is well documented to reduce Na^+^ reabsorption by PC. However, to better characterized the contribution of reabsorption and secretion pathway in WT mice under HS, we applied amiloride to inhibit a possible ENaC-mediated transport of Na^+^. As shown in Figure 4D, amiloride, as expected, has no significant effect on the JNa^+^, which is in good agreement with the decreased expression of ENaC α subunit already observed (Figure 1 and Supplemental Fig 2 and^16^). We, therefore, consider the possibility of a paracellular Na^+^ reabsorption process^14^ that would be energized by a H^+^ pump-mediated secretion of proton inducing a positive transepithelial membrane potential (Vte, as shown in CCD^14^). Indeed, the Vte measured in WT OMCD from mice under HS is 7.6±2.0 mV and decreases to 0.4±0.1 mV after bafilomycin addition (Figure 4E). Abolishment of the Vte revealed a Na^+^ secretion around –6 pmol/mm/min (Figure 4F), which is not present in the absence of the HKA2 (HKA2KO, Figure 4G). These data indicate that the secretion pathway of Na^+^, mediated by the HKA2 and NKCC1 transporters and dependent on ANP and cGMP is activated under HS diet.

**Figure 4:**
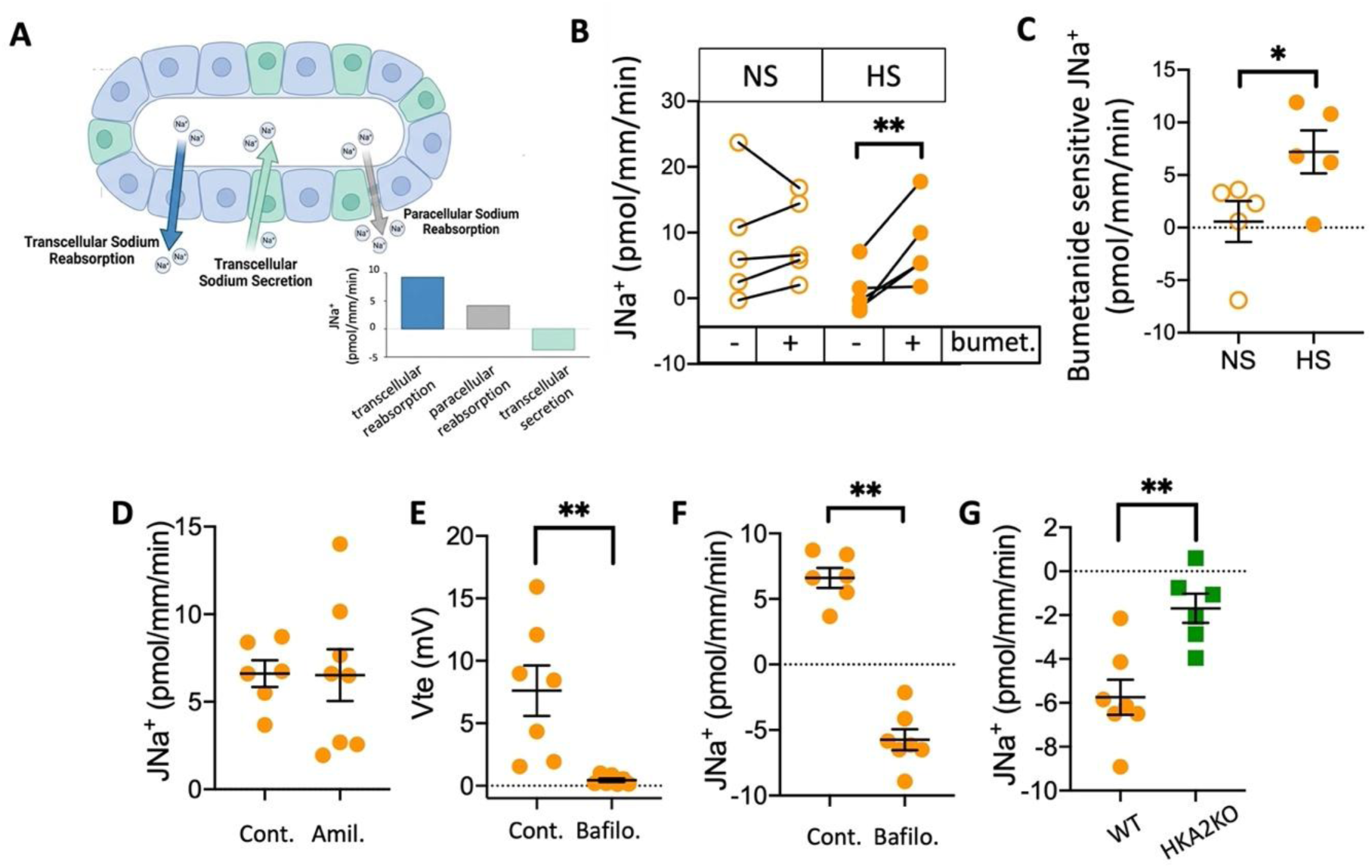
A bumetanide-sensitive and HKA2-dependent Na^+^ secretion is present in OMCD of mice fed HS diet. **A**) Schematic representation (drawn by and modified from FigureLabs) of ion fluxes in OMCD, by convention reabsorption are depicted as positive values and secretion by negative values. **B-C**) Flux of Na^+^ (JNa^+^) in microperfused isolated OMCD from WT mice under NS or HS in the absence or presence of bumetanide in the bath (basolateral side). Results are shown as individual data (**B**) and mean±s.e.m and analyzed by a paired Student t-test in **B** (** p<0.01) and by an unpaired Student t-test in **C** (* p<0.05). **D**) Flux of Na^+^ (JNa^+^) in microperfused isolated OMCD from WT mice under HS in the absence or presence of amiloride in the perfusion (lumen). Results are shown as individual data and mean±s.e.m. **E**) Transepithelial membrane potential (Vte) in microperfused isolated OMCD from WT mice under HS in the absence or presence of bafilomycin in the perfusion (lumen). Results are shown as individual data and mean±s.e.m. and analyzed by an unpaired Student t-test (** p<0.01). **F**) Flux of Na^+^ (JNa^+^) in microperfused isolated OMCD from WT mice under HS in the absence or presence of bafilomycin. Results are shown as individual data and mean±s.e.m. and analyzed by an unpaired Student t-test (** p<0.01). **G**) Flux of Na^+^ (JNa^+^) in microperfused isolated OMCD from WT and HKA2KO mice under HS in the presence of bafilomycin. Results are shown as individual data and mean±s.e.m. and analyzed by an unpaired Student t-test (** p<0.01).

### The absence of the HKA2 interferes with Na^+^ and water balance in response to high salt diet

We drew the hypothesis that the observed stimulation of the HKA2 expression in response to high salt diet would help to eliminate Na^+^ and therefore, the absence of this pump could result to Na^+^ and fluid retention and *in fine* to an increase of blood pressure. However, as shown in Figure 5A, blood pressure of HKA2KO mice is 15% lower than that of WT after 4-6 days of HS diet. In addition, we observed that under normal diet, plasma K^+^, plasma HCO_3_^−^, hematocrit and plasma Na^+^ of WT and HKA2KO mice are similar (Figure 5B-E). However, whereas WT mice maintained these parameters constant after 3 days of HS, HKA2KO mice compared to WT mice exhibited a decreased plasma K^+^ value (3.8±0.06 vs 4.35±0.15 mM, p<0.01, Figure 5B) and a higher plasma HCO_3_^−^ (27.4±0.7 vs 23.6±0.5 mM, p<0.01, Figure 5C). Moreover, HKA2KO compared to WT mice under HS diet exhibited signs of volume depletion with a higher hematocrit (40.4±0.4 vs 38.5±0.7%, p<0.05, Figure 5D) and a higher plasma Na^+^ value (155.0±1.0 vs 150.1±1.0 mM, p<0.01, Figure 5E). These results, therefore, indicate that the absence of HKA2 leads to the development of a hypovolemic, hypokalemic and alkalotic state in response to a load of salt. As shown in Figure 6A-B, WT and HKA2KO mice ingested a similar level of NS or HS food, however, if both increased their water intakes by a factor 2-3 in HS condition (Figure 6B), HKA2KO mice exhibited a higher (30-50%) consumption of water than WT mice during the whole period of HS diet. This is correlated with the increased urine volume observed in HKA2KO mice compared to WT mice (Figure 6C). Urinary Na^+^ excretion (figure 6D) is strongly increased by HS diet as expected, but we observed that the HKA2KO mice loss 20-25% more Na^+^ than WT mice (p<0.01). Regarding urinary K^+^ excretion (Figure 6E), it remained similar in WT mice whatever the diet but, in HKA2KO mice, it increased by 57% at day 1 of the HS diet (## p<0.01) and is significantly higher than in WT at day 1 and 2. The increased water intakes and water excretion by the HS diet led to a dilution of the urine in both genotypes with a higher effect on HKA2KO mice at day 2 and 3 that exhibited more diluted urine than WT mice (Figure 6F).

**Figure 5:**
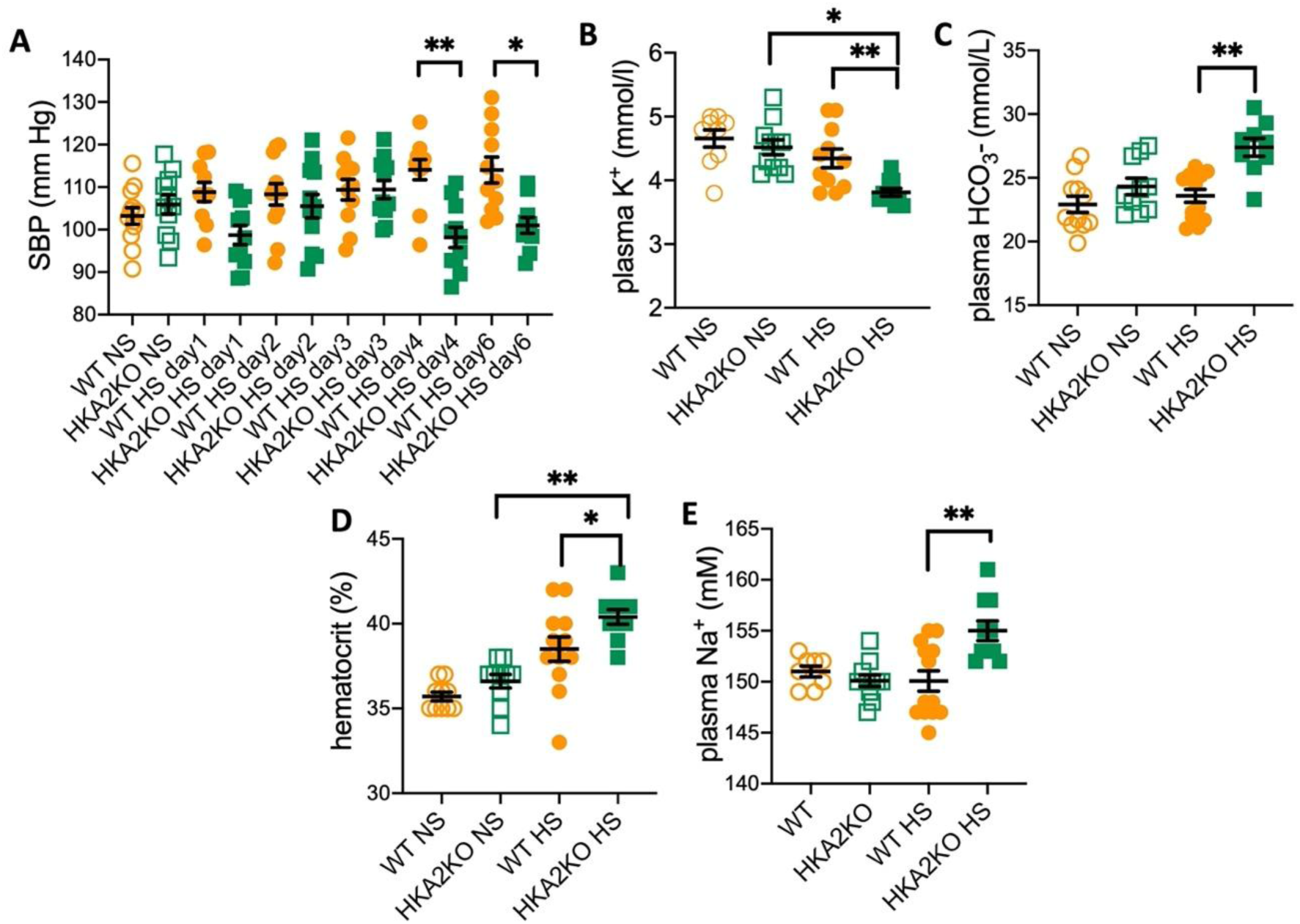
The absence of the HKA2 impacts blood pressure and plasma electrolytes concentrations in response to high salt diet. **A**) Systolic BP (SBP) was measured in WT (orange symbols) and HKA2KO (green symbols) mice submitted to a normal salt diet (open symbols) and a high Na^+^ diet (filled symbols). Results are shown as the mean±s.e.m (2 series n=11-12). Statistical analysis was performed by a two-way ANOVA test (mixed model) to measure the potential effects of genotype, time and the interaction of both and followed by a Sidak’s multiple comparison test (** p<0.01; * p<0.05) SBP is not significantly impacted by the time but depends on the genotype (p<0.01) and the interaction of both (p<0.01). **B-E**) On different series of animals (2 series, n=9-13), we measured the plasma K^+^, plasma bicarbonate, hematocrit and plasma Na^+^ levels in WT (orange symbols) and HKA2KO (green symbols) under normal salt diet (open symbols) and a high Na^+^ diet (filled symbols). Results are shown as mean±s.e.m and analyzed by a two-way ANOVA followed by a Sidak’s multiple comparison test (** p<0.01; * p<0.05). Plasma K^+^ level is not significantly impacted by the genotype of the mice but depends on the diet (p<0.01) and the interaction of both (p<0.01). Plasma bicarbonate level is neither significantly impacted by the genotype nor by the diet *per se* but by the interaction of both (p<0.01). Hematocrit level is not significantly impacted by the genotype of the mice but depends on the diet (p<0.01) and the interaction of both (p<0.01). Plasma Na^+^ level is neither significantly impacted by the genotype nor by the diet *per se* but by the interaction of both (p<0.01).

**Figure 6:**
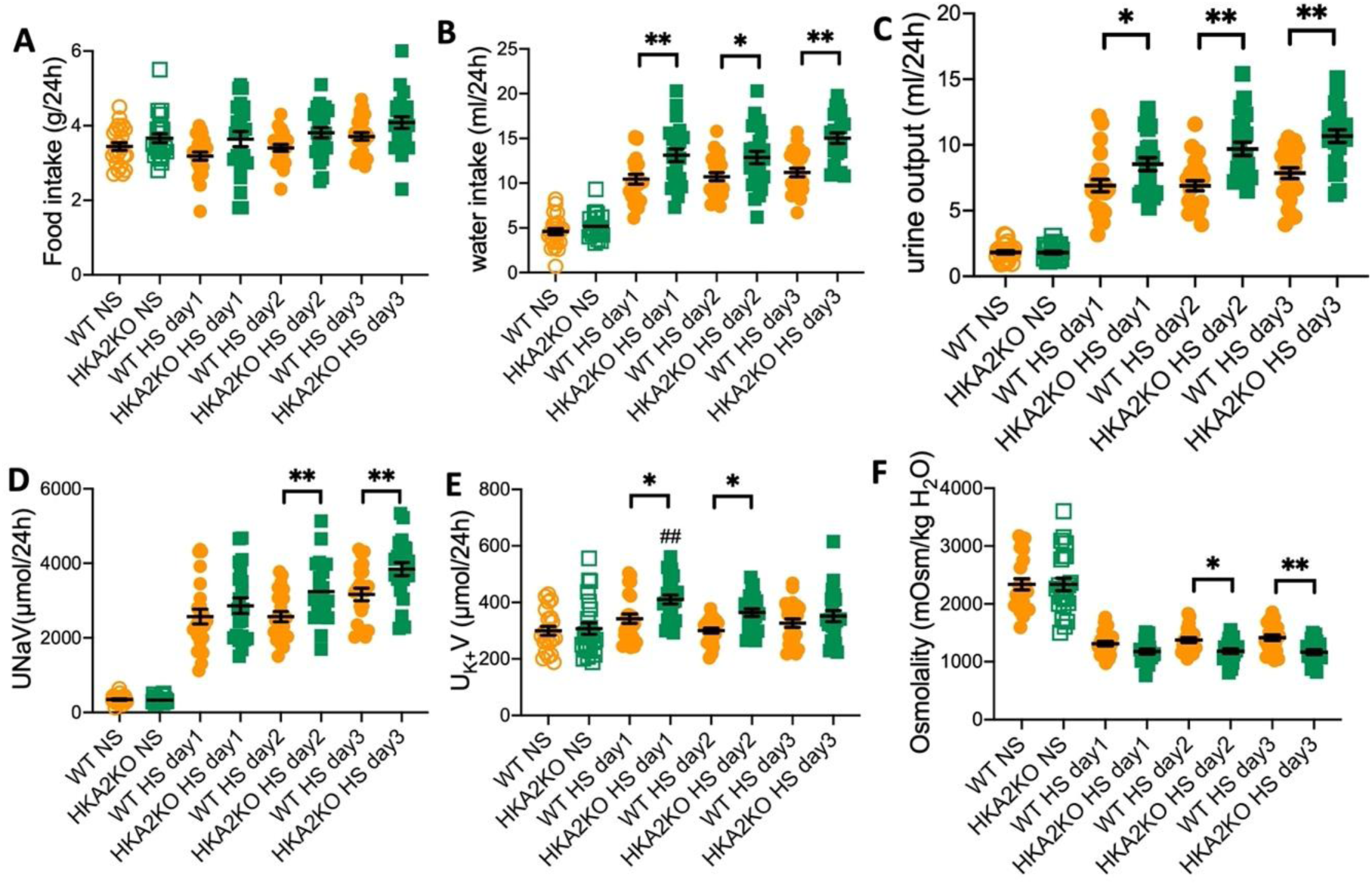
The absence of the HKA2 leads to loss of Na^+^ and fluid in response to high salt diet. **A**) Food intake (g/24h) **B**) Water intake (ml/24h) **C**) Urine output (ml/24h) **D**) Total urine Na^+^ excretion (U_Na_V, µmol/24h) **E**) Total urine K^+^ excretion (U_K_V, µmol/24h) and **F**) urine osmolality (mOsm/kg H_2_O) in WT (orange symbols) and HKA2KO (green symbols) mice submitted to a normal salt diet (NS, open symbols) and a high Na^+^ diet (HS, filled symbols). Results are shown as the mean±s.e.m (3 series, n=24-26) and analyzed by a two-way ANOVA followed by a Sidak’s multiple comparison test (WT vs KO ** p<0.01; * p<0.05 and NS vs HS day x, ## p<0.01; # p<0.05). Food intake is neither significantly impacted by the genotype of the mice, the diet type nor the interaction of both. Water intake is significantly impacted by the genotype (p<0.01), by the diet (p<0.01) but not by the interaction of both. Urine output is significantly impacted by the genotype (p<0.01), by the diet (p<0.01) and by the interaction of both (p<0.01). Urine Na^+^ excretion is significantly impacted by the genotype (p<0.01), by the diet (p<0.01) but not by the interaction of both. Urine K^+^ excretion is significantly impacted by the genotype (p<0.01), by the diet (p<0.01) and by the interaction of both (p<0.05). Urine osmolality is significantly impacted by the genotype (p<0.01), by the diet (p<0.01) but not by the interaction of both.

### The absence of HKA2 shuts down the Na^+^ reabsorption in the thick ascending limb

To understand the mechanism involved in the loss of fluid and Na^+^ in HKA2KO mice under HS diet, we performed acute pharmacological treatments. As shown in Figure 7A, under NS diet, furosemide had a similar effect on Na^+^ excretion in WT and HKA2KO mice where it induced a 6-10-fold increase of Na^+^ excretion. Under HS diet, furosemide increased Na^+^ excretion by only a factor 2 in WT, indicating that NKCC2 *in vivo* activity is reduced but not completely inhibited. Conversely, in HKA2KO mice, furosemide had no significant effect on Na^+^ excretion indicating that NKCC2 *in vivo* activity is turned off in HS diet. As shown in Figure 7B, under NS diet, hydrochlorothiazide (HCTZ) had a similar effect on Na^+^ excretion in WT and HKA2KO mice where it induced a 3-fold increase of Na^+^ excretion. Under HS diet, HCTZ did not increase significantly Na^+^ excretion in WT or HKA2KO mice, indicating that NCC *in vivo* activity is strongly reduced by HS diet. This is supported by the similar decrease of pNCC expression under HS in both WT and HKA2KO mice (Supplemental Figure 2), indicating that the K^+^ switch process leading to NCC stimulation in hypokalemic situation is blunted in a context of HS and/or in the absence of HKA2 as shown previously^14^. As shown in Figure 7C, under NS diet, amiloride had a similar effect on Na^+^ excretion in WT and HKA2KO mice where it induced a 4-5-fold increase of Na^+^ excretion. Under HS diet, amiloride did not increase significantly Na^+^ excretion in WT or HKA2KO mice, indicating that ENaC *in vivo* activity is strongly reduced by HS diet as directly observed by microperfusion analysis. To investigate further the difference of NKCC2 *in vivo* activity between WT and HKA2KO mice under HS, we analyzed NKCC2 protein expression. Western blot analysis on renal membrane preparations targeting NKCC2 and its phosphorylated form (Figure 7D) and observed that NKCC2 and pNKCC2 expression were reduced in HKA2KO mice under HS compared to WT mice. A possible consequence of a decreased activity of NKCC2 in the TAL is a decrease of the GFR^17^. As shown in Figure 7E, transdermal monitoring on conscious animals revealed that HS did not modify GFR in WT (974 vs 910 µl/min/100g) but decreased it by 30% in HKA2KO mice (992 vs 714 µl/min/100g, p<0.05).

**Figure 7:**
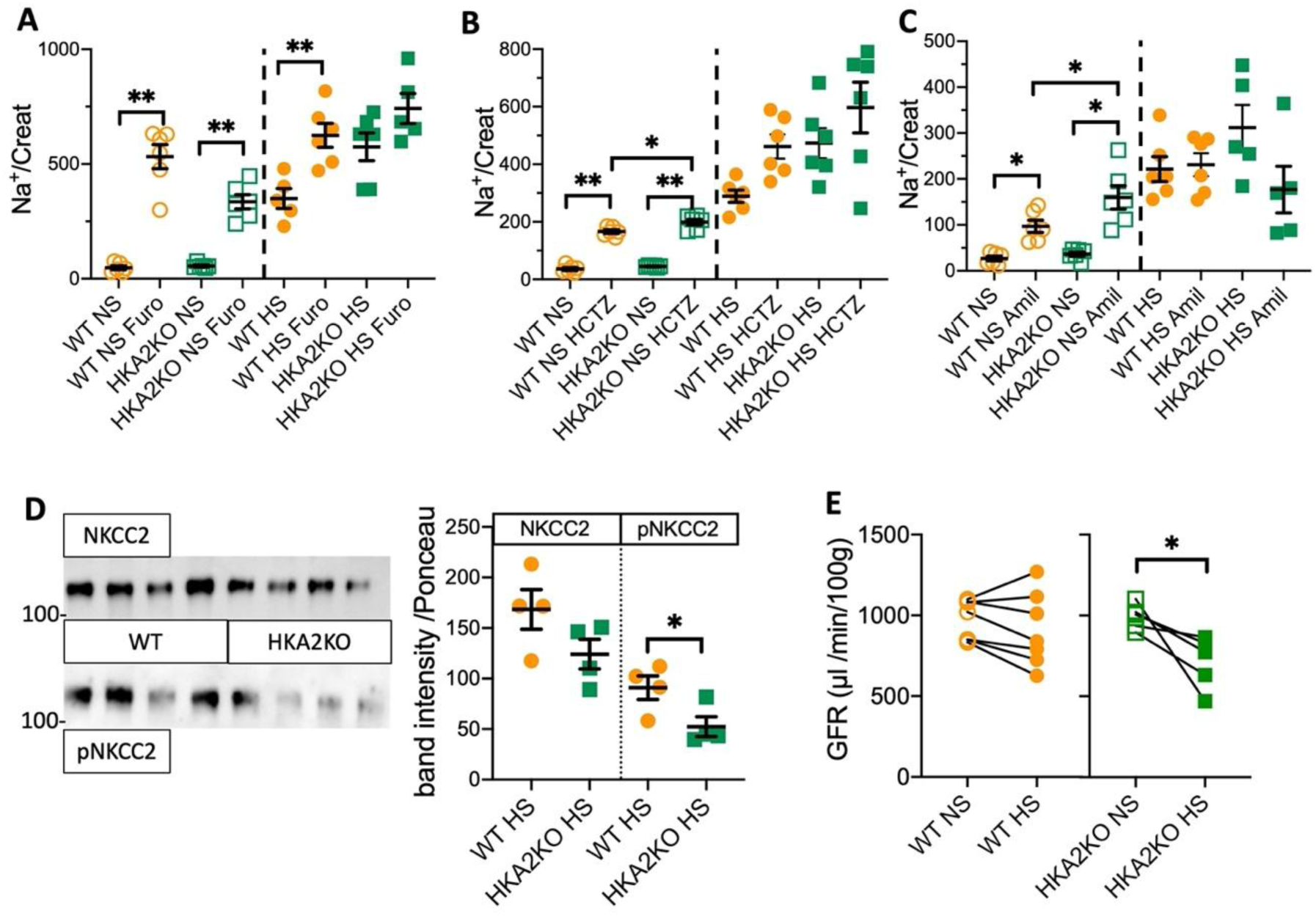
Na^+^ reabsorption by NKCC2 in the thick ascending limb is strongly reduced in the absence of the HKA2 in response to high salt diet. Acute natriuretic response to furosemide (**A**), hydrochlorothiazide (HCTZ, **B**) and amiloride (**C**) in WT (orange symbols) and HKA2KO (green symbols) mice submitted to a normal salt diet (NS, open symbols) and a high Na^+^ diet (HS, filled symbols). Results are shown as the mean±s.e.m (n=6) and analyzed by a one-way ANOVA followed by a Sidak’s multiple comparison test (** p<0.01; * p<0.05). **D**) Protein expression of NKCC2 and phosphoNKCC2 and the quantification of the band intensities normalized by the red Ponceau labelling. Results are shown as the mean±s.e.m (n=4) and analyzed by a Student t-test (* p<0.05). **E**/ GFR measurements on the same animals under normal salt diet (NS) or after 3 days of high salt diet (HS). Results are shown as the mean±s.e.m (n=7 for WT and 5 for HKA2KO mice) and analyzed by a paired Student t-test (* p<0.05).

### The absence of HKA2 induces vasopressin resistance

As shown in Figure 6C, the HKA2KO mice under HS exhibited a more important loss of fluid than WT mice. To investigate further this aspect, we measured 24h urine copeptin secretion (a marker of arginine-vasopressin peptide; AVP) and showed that it increased in both genotypes under HS diet (Figure 8A). However, the HKA2KO mice exhibited a 2-time higher urine copeptin excretion than the WT mice at day 1 and 2 of the HS diet, indicating that the HKA2KO are likely in a state of dehydration sooner than WT mice. To test for the V2R-dependent-response to AVP, we compared the urine osmolality in WT and HKA2KO mice before and after a single injection of ddAVP. The ddAVP effect (ratio urine osmolality after/before ddAVP injection) in WT mice is not significantly changed by the HS diet compared to the NS diet (Figure 8B). However, whereas ddAVP induced a 50-60% increase of urine osmolality in HKA2KO mice under NS diet, ddAVP effect is significantly (p<0.05) lower under HS diet, indicating that in this condition the activation of the V2 receptor-dependent pathway is blunted and HKA2KO mice could be considered as “resistant” to ddAVP (Figure 8C) in terms of urine concentration. The analysis of AQP2 protein expression confirmed this result, showing that AQP2 is 3-fold increased by HS diet in WT mice but not in HKA2KO mice (Figure 8D). Interestingly, by acting on V1a receptor, AVP also stimulates the intrarenal production of prostaglandins such as PGE2 that contribute to Na^+^ and water excretion by antagonizing the effect of AVP on the V2 receptor^18^. As shown in Figure 8E, the renal expression of Cox-2, a key enzyme of the prostaglandins synthesis is upregulated in HKA2KO mice under HS compare to WT under HS. In parallel, the urine excretion of PGE2 is significantly, rapidly and steadily increased (by 100% compared to the NS condition) after HS. Conversely, PGE2 excretion in WT mice under HS remained similar to that in NS condition during the first two days of the challenge. It is significantly increased by 80% at the third day of the challenge. These results suggest that a vicious circle is going on in HKA2KO mice where loss of water and Na^+^ induce AVP production, which in turns, stimulates PGE2 production that would antagonize the V2 receptor-dependent antidiuretic and anti-natriuretic effect of AVP, resulting, *in fine,* to an increased loss of fluid and salt.

**Figure 8:**
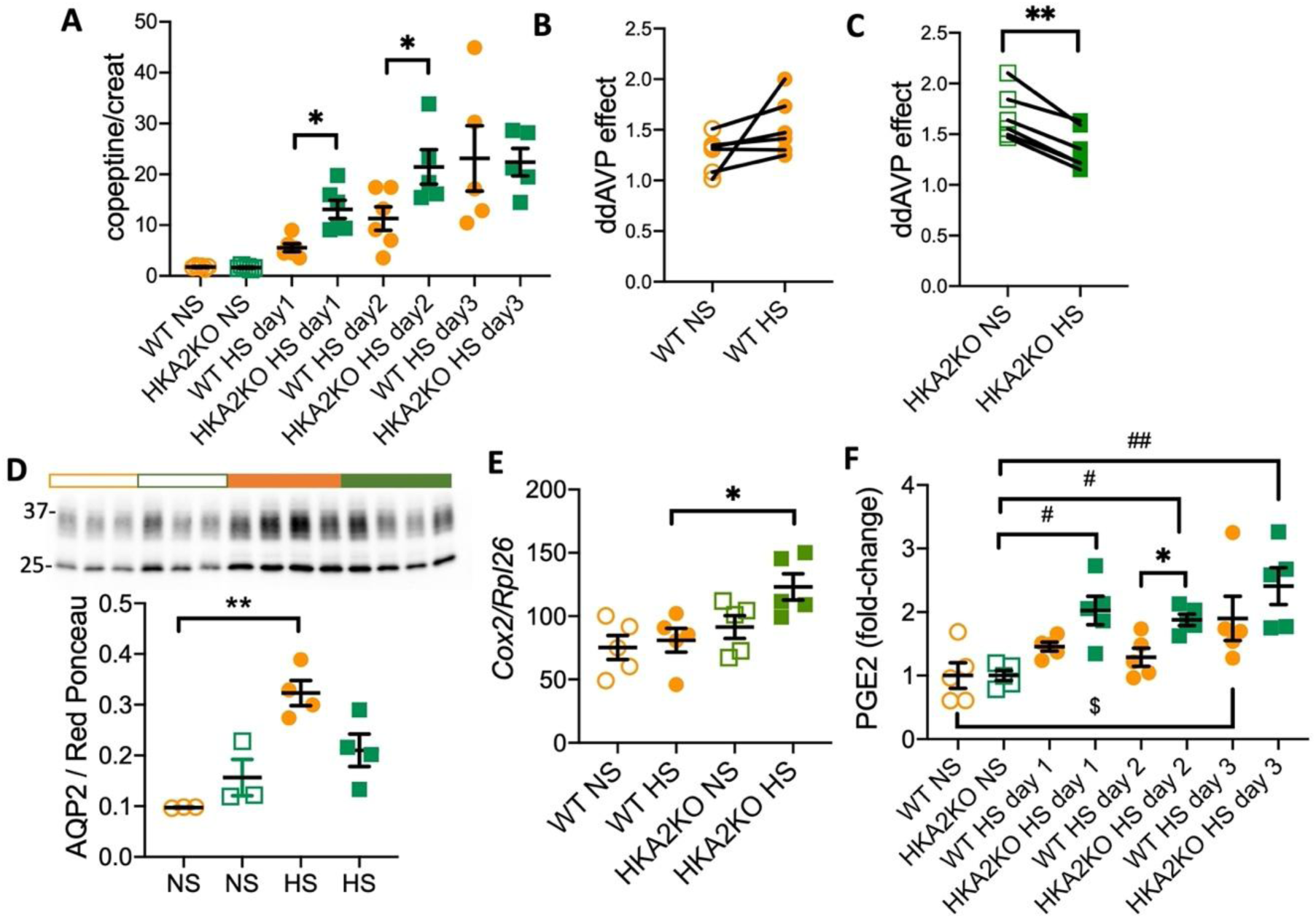
Vasopressin resistance in the absence of HKA2 in response to high salt diet. **A)** Urine copeptine excretion in WT (orange symbols) and HKA2KO (green symbols) mice submitted to a normal salt diet (NS, open symbols) and a high Na^+^ diet (HS, filled symbols). Results are shown as the mean±s.e.m (n=5-6) and analyzed by a two-way ANOVA followed by a Sidak’s multiple comparison test (WT vs KO * p<0.05). Urine copeptine is significantly impacted by the genotype (p<0.01), by the diet (p<0.01) but not by the interaction of both. **B-C**) Urine osmolality measured after ddAVP injection normalized by those obtained with the same animal before the injection in mice under normal salt diet (NS) or after 3 days of high salt diet (HS). Results are shown as the mean±s.e.m (n=6) and analyzed by a paired Student t-test (** p<0.01). **D)** Protein expression of AQP2 and the quantification of the band intensities normalized by the red Ponceau labelling. Results are shown as the mean±s.e.m (n=3 in NS and 4 in HS) and analyzed by a one-way ANOVA (** p<0.01). **E)** *Cox2* expression in total kidney (n=5) normalized by the housekeeping gene *Rpl26* in WT or in HKA2KO mice under NS or HS diets. Results are shown as mean±s.e.m and analyzed by an unpaired Student t-test (*p<0.05). **F**) Urinary PGE2 (n=5) in WT or in HKA2KO mice under NS or HS diets normalized by the mean value of the PGE2 excretion (pg/24h) of WT (596±120 pg/24h) or HKA2KO mice (481±42 pg/24h). Results are shown as the mean±s.e.m and analyzed by a two-way ANOVA followed by a Sidak’s multiple comparison test (WT vs KO * p<0.05 and NS vs HS day x, ## p<0.01; # p<0.05).

## Discussion

In this study, we propose that renal adaptation to a salt load involves the secretion of Na^+^ through the AIC of the collecting duct in addition to the inhibition of Na^+^ reabsorption along the nephron^19^. The secretion of Na^+^ into the lumen at the end of the collecting duct is essential to prevent over-inhibition of Na^+^ reabsorption in the TAL, thereby maintaining the balance of Na^+^, K^+^, and water.

The role of AIC in responding to a high-salt diet is demonstrated by the increased number of these cells, a characteristic seen in conditions where their transport properties are crucial. AICs are well-known for increasing their number during acidosis^20^ through proliferation^12,21,22^ or transdifferentiation^23^. A similar increase (by 10-15 %) is observed with dietary K^+^ restriction^13,24,25^. We favor the hypothesis of a proliferation mechanism instead of a transdifferentiation process since 1/ RNAseq analysis shows up-regulation of pathways linked to proliferation, confirmed by increased Pcna expression and 2/ transdifferentiation processes, particularly from B-intercalated cells (BIC) to AIC, are unlikely as the medullary collecting duct lacks BIC. Moreover, the increase in total cell number per mm likely reflects additional cells rather than replacement.

For decades, the renal control of Na^+^ balance focused on reabsorption processes, involving their stimulation or inhibition. However, evidence for Na^+^ and Cl^−^ secretion in the collecting duct exist in the literature. In the mid-80’s^26^, measurements of unidirectional Na^+^ fluxes (from bath to lumen i.e. secretion and lumen to bath i.e. reabsorption) in CCD from rat under normal condition showed that there are equivalent Na^+^ fluxes in both direction, resulting in a null net flux. Treatment with DOCA salt or AVP only stimulated the reabsorption of Na^+^ without affecting the Na^+^ secretion. In 1990, Rocha and Kudo^27^ showed that Cl^−^ secretion depended on ANP and was sensitive to furosemide when applied at the basolateral side of the cells. Small but significant Na^+^ and Cl^−^ secretion in the CCD of mice with inhibited Na^+^ reabsorption pathways has been reported by Chambrey et al.^8,28^. In 2012, Wall’s group demonstrated Cl^−^secretion through AIC mediated by NKCC1 under forced Na^+^ reabsorption (aldosterone infusion by minipump) or high Na^+^ diet conditions^29^. Since the activation of NKCC1 promotes also the entry of Na^+^ into the AIC, well-known to not express the Na,K-ATPase, the fate of this cation that cannot accumulate into the intracellular space was questioned. In 2016, we showed that the HKA2 acts as an apical Na,K-ATPase transporting Na^+^ out of the cell^14^. Indeed, this pump exhibits features of both H,K– and Na,K-ATPase (for review see^30^) and can transport Na^+^ with the same affinity than the Na,K-ATPase^31–33^. The ion specificity of the HKA2 (Na^+^ *vs* H^+^) was controlled by the ANP/cGMP hormonal system^15^. ATP12a down-regulation in hypertensive rats^34^ and its identification in independent GWAS studies investigating systolic and diastolic blood pressure related genes^35–38^, in addition to our present data, likely confirms that the HKA2 is involved in the Na^+^ balance and blood pressure regulation by promoting Na^+^ secretion.

The absence of this pathway should impede proper Na^+^ secretion, leading to Na^+^ retention and eventually increased blood pressure. Surprisingly, we observed the exact opposite, as the HKA2KO mice under HS lost more Na^+^ and fluid and had lower blood pressure. This confirms that the HKA2 plays a role in the salt and water balances but also suggests that a mechanism is triggered to compensate for the lack of Na^+^ secretion. This mechanism consists in an exaggerated anti-natriuretic response mediated by an over production of PGE2 that would inhibit NKCC2^39^ and blunt the effects of AVP mediated by its V2 receptor^40^. Therefore, to compensate for the absence of Na^+^ secretion, that would permit to finely tune the total renal Na^+^ excretion with the intakes, the kidney decreases its capacity of Na^+^ reabsorption by strongly inhibiting a major Na^+^ reabsorption pathway, leading, finally to a loss of Na^+^. If it may seem curious to observe an exaggerated loss of salt in response to a high salt diet, we note that it has been already observed, in hepsin KO mice, where uromodulin aggregation leads to tubular damages and TAL dysfunction^41^.

The description of a novel pathway stimulating Na^+^ secretion in the kidney in response to salt loading is of major importance and represents a significant paradigm shift in comparison with the current dogma. The Na^+^ balance, at the kidney level, is therefore control by both reabsorption and secretion processes.

## Acknowledgements

Physiological studies have been performed with the help of Gaëlle Brideau and Nadia Frachon from the “plateforme d’exploration fonctionnelle du petit animal” of the team “Physiologie Rénale and Tubulopathies” at the Centre de Recherche des Cordeliers. We are grateful for the technical assistance of the Centre d’Exploration Fonctionelle crews in the management of our colonies of mice. This study was supported by the Agence National de la Recherche (ANR) project ANR-21-CE14-0040-01 (GC), The Fondation pour La Recherche Médicale (FRM) project DPC20171138949 (GC), the Fondation Lefoulon Delalande (CR), the Prolific association (CR) and the Société Francophone de Néphrologie, Dialyse et Transplantation (SFNDT) project R20186DD, CR).

---

**Supplemental Figure 1:**
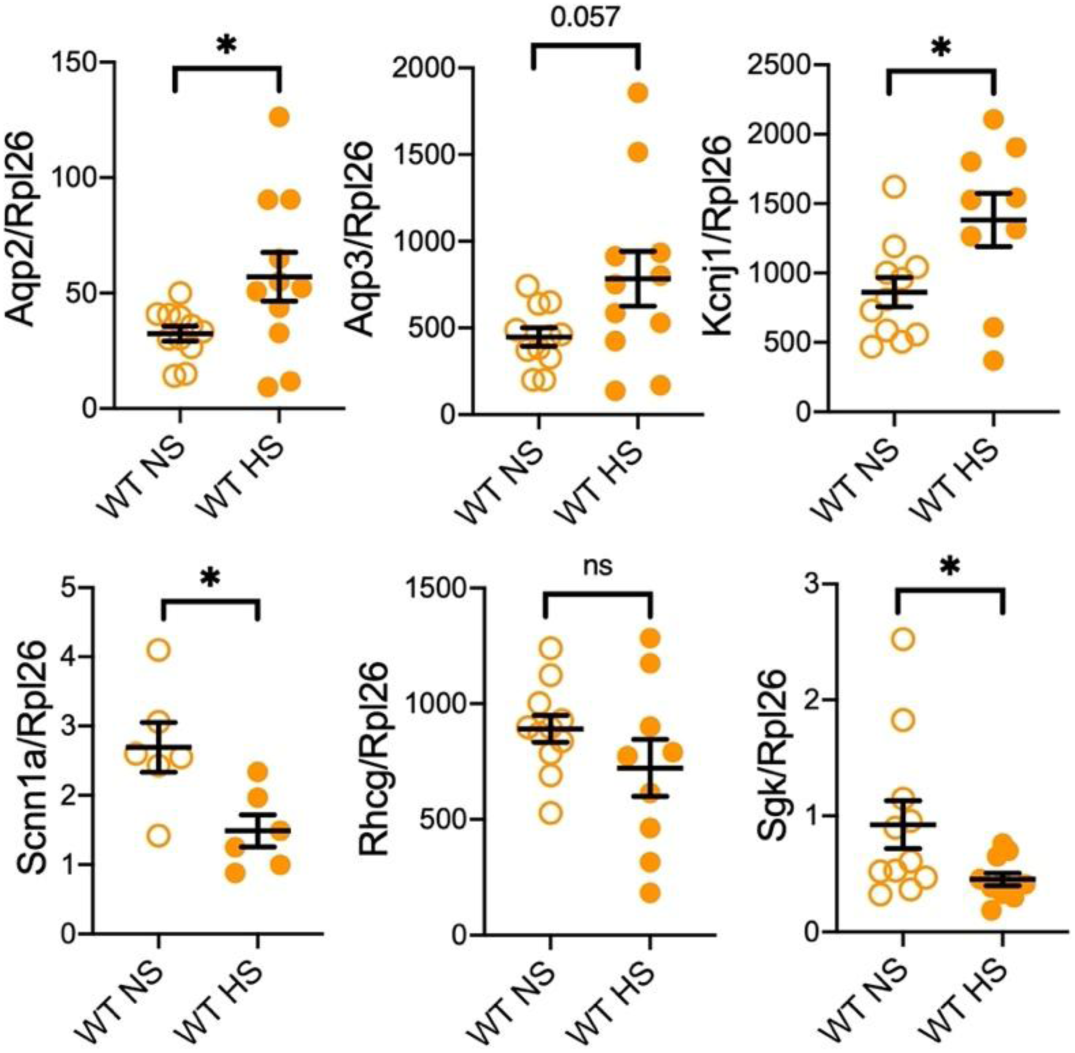
Modification of gene expression in collecting ducts of wild-type mice. Expression of some up– and downregulated genes found in the RNAseq in isolated OMCD (n=7-11) normalized by the housekeeping gene *Rpl26* in WT mice under NS or HS diets. Results are shown as mean±s.e.m and analyzed by an unpaired Student t-test (*p<0.05, ** p<0.01).

**Supplemental Figure 2:**
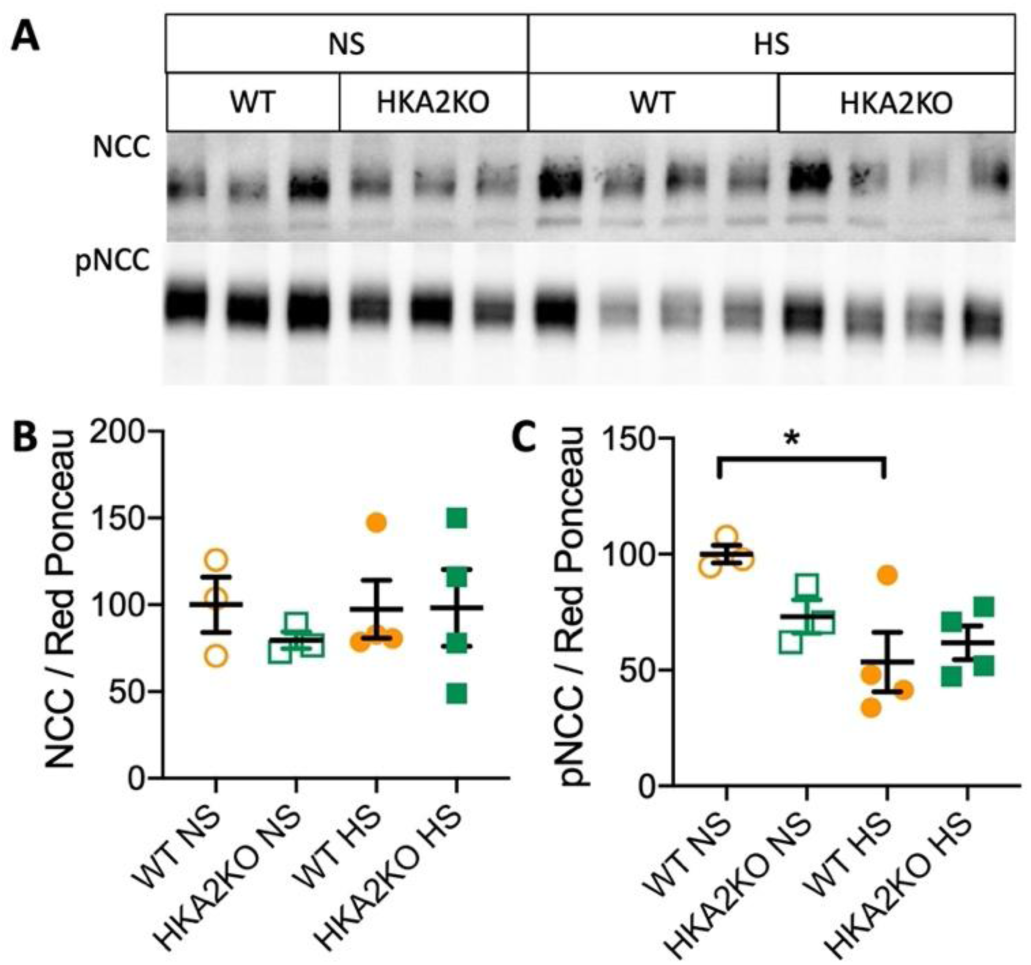
**A**) Protein expression of NCC and phosphoNCC. **B-C)** Quantification of the band intensities normalized by the red Ponceau labelling. Results are shown as the mean±s.e.m (n=3 for NS diet and 4 for HS diet) and analyzed by a Student t-test (* p<0.05).

**Supplemental Figure 3:**
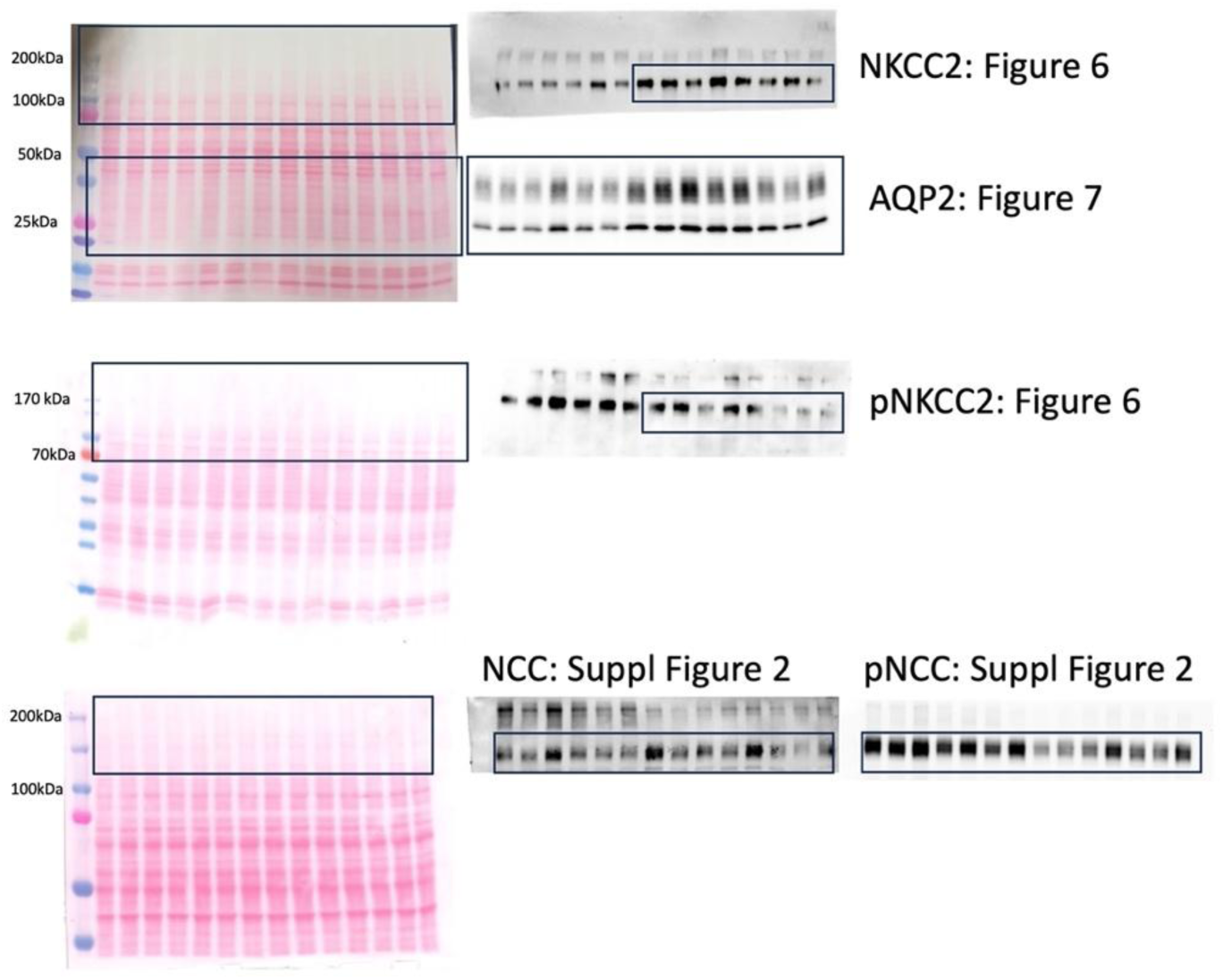
Full Western blots and red Ponceau staining.

