## Supplemental Material and Methods for "Renal adaptation to high salt diet requires tubular Na^+^ secretion through type A intercalated cells"

**Metabolic analysis**

Urinary creatinine concentrations were determined using an automatic analyzer (Konelab 20i; Thermo, Cergy Pontoise, France). Urinary $Na^+$ and $K^+$ concentration was determined by flame photometry (M420, Sherwood Scientific, France). Osmolality was measured on 24h-urine collection using an osmometer (Vogel 6300 Voebling osmometer, Bioblock Scientific). Plasma parameters were measured by retro-orbital puncture on the anesthetized animal (a mix of xylazine, 10mg/kg and ketamine 100mg/kg) with an Epoc pH/blood-gas analyzer (Siemens Healthineers, Saint Denis, France). Glomerular filtration rate was determined using transcutaneous monitoring of the decrease in fluorescence after intravascular injection of FITC-Sinistrin (batch VE17202021, Fresenius Kabi, Linz, Austria). The fluorescence detector, a miniaturized photodiode (MediBeacon GmbH, Mannheim, Germany), was attached to the left flank of the animal previously depilated and protected from any light source by placing occulting paper. A dose of 8 mg/100 g of body weight (concentration of 20 mg/ml) of Fluorescein-Isothiocyanate (FITC)-Sinistrin was injected via femoral artery 10 min after placing the camera to obtain a baseline value. The fluorescence peak was then obtained approximately 10 min after injection of FITC-Sinistrin. The camera measured the transcutaneous fluorescence emitted from the dermal capillaries over a total period of 2 h. The GFR was calculated by the MB Lab/MB Studio software (MediBeacon GmbH, Mannheim, Germany) from the slope of the decrease in fluorescence intensity over time according to a tri-compartmental model with linear regression

Measurement of urinary copeptine (Clinisciences, #CEA365Mu-96), cGMP (Abcam, ab65356), PGE2 (Cayman Chemical #514010) and of plasma ANP (Sigma-Aldrich, RAB00385) were performed using ELISA kit according to the supplier's informations.

**mRNA and protein expression analysis**

RNAs were extracted from whole kidneys using the TRI reagent (Invitrogen, France) following the manufacturer's instructions. One µg of total RNA was then reverse-transcribed (Roche Diagnostics, France) and real-time PCRs were performed on a LightCycler (Roche Diagnostics, France) as previously described [1].

Total kidneys were homogenized in a lysis buffer (250 mM sucrose, 100 mM Tris-Hepes, pH 7.4 and protease and phosphatase inhibitors cocktail (Complete, Roche Diagnostics)). After removal of aggregates and nuclear-associated membrane by low-speed centrifugations, the plasma membrane enriched fraction (17000 x g for 30 min) was recovered into the lysis buffer and its protein content was measured with the BCA method (Thermo Scientific Pierce). Forty µg of protein were then denaturated, resolved by SDS-PAGE (10% polyacrylamide) and transferred onto a nitrocellulose membrane. Ponceau red labelling was carried out to check for protein loading accuracy. Western blots were performed according to the standard procedure using a rabbit anti-NKCC2 (Merck Millipore, 1/1000), a rabbit anti-pNKCC2Ser91 (Dundee, 1/500) and a goat-anti-AQP2 (Santa Cruz). For quantification, the band intensities were normalized by Ponceau red intensity.

**AIC isolation**

Anesthetized mice underwent perfusion in the right ventricle with a digestion solution composed of Krebs buffer containing Liberase (Roche). Subsequently, both kidneys were decapsulated, minced with razor blades, and incubated in the digestion solution for 22 minutes at 37°C. The tube was agitated thoroughly every 5 minutes for 5 seconds to disperse kidney tubules and cells. After 22 minutes, the reaction was halted at 4°C. The tubular and cell suspension underwent sequential filtration through 212µm, 100µm, and 70µm sieves. The flowthrough was centrifuged at 4°C for 10 minutes. The pellet was resuspended in ice-cold Krebs and centrifuged again to eliminate debris. Remaining erythrocytes were lysed with Red Blood Cell Lysis buffer (Thermo Scientific). Cells were resuspended in Krebs buffer with 0.05% BSA and 1mM EDTA. APC anti-mouse CD117 (Biolegend) and PE/Cy7 anti-mouse CD45 (Biolegend) were employed to label the cells. CD45-CD117+ cells were sorted using a cell sorter, FACS ARIA II (BD Biosciences). Live/Dead staining kits were utilized to include only living cells during sorting, with a viability of 90% for CD45-CD117+ cells.

**Sodium flux in isolated microperfused OMCD**

Concentrations of $Na^+$ and creatinine were determined by HPLC using 14 nl of collected fluid. For each collection period, the flux of ion X per unit length of tubule ($J_X$) was calculated as $J_X = ([X]_p \times V_p - [X]_c \times V_c)/(L \times t)$, where $[X]_p$ and $[X]_c$ are the concentrations of ion $X$ in the perfusate and collectate, respectively; $V_p$ and $V_c$ are the perfusion and collection rates, respectively; $L$ is the tubule length; and $t$ is the collection time. $V_p$ was calculated as $V_p = V_c \times [\text{creat}]_c/[\text{creat}]_p$, where $[\text{creat}]_c$ and $[\text{creat}]_p$ are the concentrations of creatinine in the collectate and perfusate, respectively. For each tubule, and each period, ion fluxes were averaged over the collections of a period.

1. Grimont A, Bloch-Faure M, El Abida B, Crambert G. Mapping of sex hormone receptors and their modulators along the nephron of male and female mice. *FEBS Lett*. May 19 2009;583(10):1644-1648.
